# MLL-AF9 Initiates Transformation From Fast-Proliferating Myeloid Progenitors

**DOI:** 10.1101/484220

**Authors:** Xinyue Chen, Amaleah Hartman, Xiao Hu, Anna E. Eastman, Xujun Wang, Mei Zhong, Shangqin Guo

## Abstract

Cancer is a hyper-proliferative clonal disease. Whether the proliferative state originates from the cell-of-origin or emerges later remains elusive. By tracking *de novo* transformation from normal hematopoietic progenitors expressing an acute myeloid leukemia (AML) oncogene MLL-AF9, we reveal that the cell cycle rate heterogeneity among granulocyte-macrophage progenitors (GMPs) determines their probability of transformation. An intrinsic fast cell cycle kinetics at the time of oncogene expression provide permissiveness for transformation, with the fastest cycling 3% of GMPs (∼0.006% of bone marrow nucleated cells) acquiring malignancy with nearly 100% efficiency. Molecularly, we propose that MLL-AF9 preserves the gene expression of the cellular states in which it is expressed. As such, when expressed in the naturally-existing, rapidly-cycling myeloid progenitors, this cell state is perpetuated, yielding malignancy. Our work elucidates one of the earliest steps toward malignancy and suggests that modifying the cycling state of the cell-of-origin could be an effective approach to prevent malignancy.

---

Not all cells respond to oncogenic insults which result in malignant transformation^1^, raising the question of what discriminates a cell to be transformed from those to remain normal despite their underlying genetic abnormalities. A frequently considered scenario is the acquisition of additional genetic lesions, allowing mutant cells to gain proliferative advantage, evade apoptosis and/or immune surveillance^2^, leading to their net expansion. While this multi-hit oncogenic model has extensive support from solid cancers^3–5^, several types of malignancy have rather low mutational load, such as those of hematopoietic origin^5,6^. Expression of a single oncogene, such as the MLL fusion oncogenes, is often sufficient to induce malignancy in animal models^7–9^. Alternatively, different cells could depend on distinct gene products so that specific oncogenes only affect selected cell types, as seen in the early development of retinoblastomas when Rb is lost^10^, or in breast and ovarian cancers when BRCA1 is mutated^11^. But, even when present in the relevant target cell types, oncogenes may not lead to immediate transformation. For example, the chronic myeloid leukemia (CML) driver BCR-ABL can persist in hematopoietic stem cells (HSCs) without causing aggressive malignancy^12^. Furthermore, it is conceivable that oncogenic mutations only give rise to malignancy when acquired by rare stem cells. However, when malignancy is manifested by progeny of the mutated stem cells, it is difficult to ascertain whether transformation is initiated in the stem cells themselves or their differentiated descendants. Indeed, stem cells could even resist transformation as compared to their more differentiated descendents^13^. Overall, the acquisition of malignancy appears to follow yet unappreciated rules.

In this report, we set out to determine the cellular traits that contribute to the acquisition of *de novo* malignancy. Specifically, we focused on granulocyte-macrophage progenitors (GMPs), which are permissive for MLL-fusion oncogene mediated transformation^7,8^. GMPs expressing an MLL-fusion oncogene could produce two types of progeny: differentiated ones despite the oncogene expression, or malignant ones that could eventually develop into lethal AML *in vivo*. This binary system provides a unique opportunity to dissect the molecular and cellular differences that help to drive malignancy.

## Tracking single hematopoietic cells from normal to malignant

To unveil potential principles governing the emergence of malignancy, we used an AML model, for which a single oncogene MLL-AF9 is sufficient to initiate a lethal disease^7,8^. An ideal experimental system to discern cellular traits underlying the permissiveness to transformation should satisfy the following criteria: (1) all cells are similar in developmental stage and oncogene expression, but only some transform to a malignant state; (2) the normal cellular behavior independent of oncogene effects can be assessed and (3) related to whether that cell transforms.

To achieve controlled oncogene expression, we generated an inducible MLL-AF9 (iMLL-AF9) allele, in which cDNA encoding human MLL-AF9 oncogene followed by an IRES-**Δ**NGFR cassette^14^, was targeted into the *Hprt* locus under the control of a tetracycline response element (TRE)^15^. This allele was crossed with a constitutively expressed reverse tetracycline transactivator (rtTA) allele^16^ (Fig. 1a) to enable doxycycline (Dox) inducible MLL-AF9 expression, which could be monitored by the co-expressed **Δ**NGFR on cell surface (Extended Data Fig. 1a). The iMLL-AF9 allele eliminates variability in oncogene copy number or integration sites introduced via viral transduction^7,8,14^. Further, precisely timed oncogene induction by Dox addition enables cellular states assessment before and after oncogene induction.

**Fig 1:**
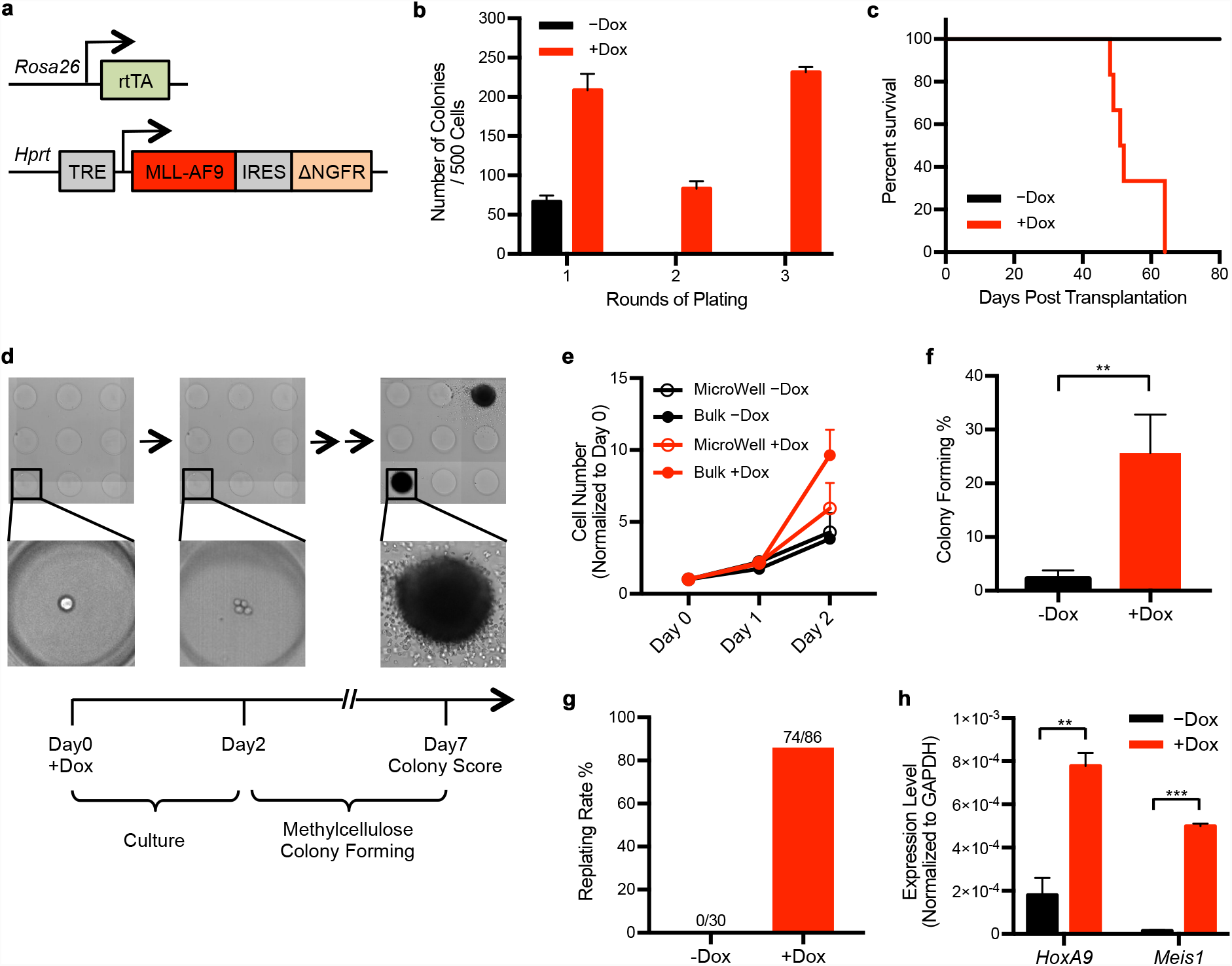
Tracking MLL-AF9 mediated transformation from single hematopoietic cells. a, Schema of the inducible MLL-AF9-ires-NGFR allele targeted into the endogenous *Hprt* locus. b, Dox-dependent serial colony formation by iMLL-AF9 GMPs. n=3 for both conditions, p<0.001 for all comparisons. c, Kaplan-Meier curve of cohorts of mice transplanted with 12,000 iMLL-AF9 GMPs, fed with control or Dox water (n=6 for each group). AML pathology was confirmed and shown in Extended Data Fig. 1b-e. d, Representative images tracking single GMPs and their progeny in micro-wells. Cells in micro-wells were kept in liquid culture for two days, after which the medium was replaced by methylcellulose (MethoCult GF M3434) to allow colony formation. The presence of colonies was verified after another 5 days (for a total of 7 days in culture). e, Comparison of iMLL-AF9 GMP proliferation rates in micro-wells and in bulk culture. The proliferation rates in mi cro-wells were calculated by adding up all cell numbers from individual wells. f, Colony forming efficiencies from single iMLL-AF9 GMPs following the schema in d. The presence of single cells was confirmed at day 0. n=4 for -Dox, n=3 for +Dox, p=0.0012 from unpaired t-test. g, Single colonies, as shown in d, were plucked and re-plated in methylcellulose for secondary colony formation. The percentages of colonies that gave rise to replatable colonies are plotted. None of the 30 colonies formed in the absence of Dox supported replating, while 74 of the 86 colonies formed in the presence of Dox supported replating. h, RT-QPCR analyses of *HoxA9* and *Meis1* in colonies formed by single iMLL-AF9 GMPs +/-Dox. p=0.002 for *HoxA9,* p<0,001 for *Meis1.*

We detected transformation based on serial colony forming capacity in methylcellulose, a well-accepted surrogate assay for hematopoietic malignancies^9^, and validated the pathology *in vivo*. As expected, iMLL-AF9 GMPs led to serial colony formation *in vitro* (Fig. 1b) and lethal AML *in vivo* in a Dox dependent manner (Fig. 1c, Extended Data Fig. 1b-e), confirming the transformability of GMPs. However, bulk cultures obscure cellular heterogeneity with regard to transformation permissiveness. We therefore plated single iMLL-AF9 GMPs in micro-wells to visualize individual cells as well as their progeny (Fig. 1d, Extended Data Fig. 2a-c). Cells in micro-wells proliferated at a comparable rate to those in bulk cultures (Fig. 1e) and were capable of forming compact colonies when methylcellulose was added (Fig. 1d). To confirm transformation, primary colonies arising from single iMLL-AF9 GMPs in micro-wells were plucked and re-plated into 96-well plates for secondary colony formation (Extended Data Fig. 2a). The presence of secondary colonies was scored as an event of malignant transformation.

**Fig 2:**
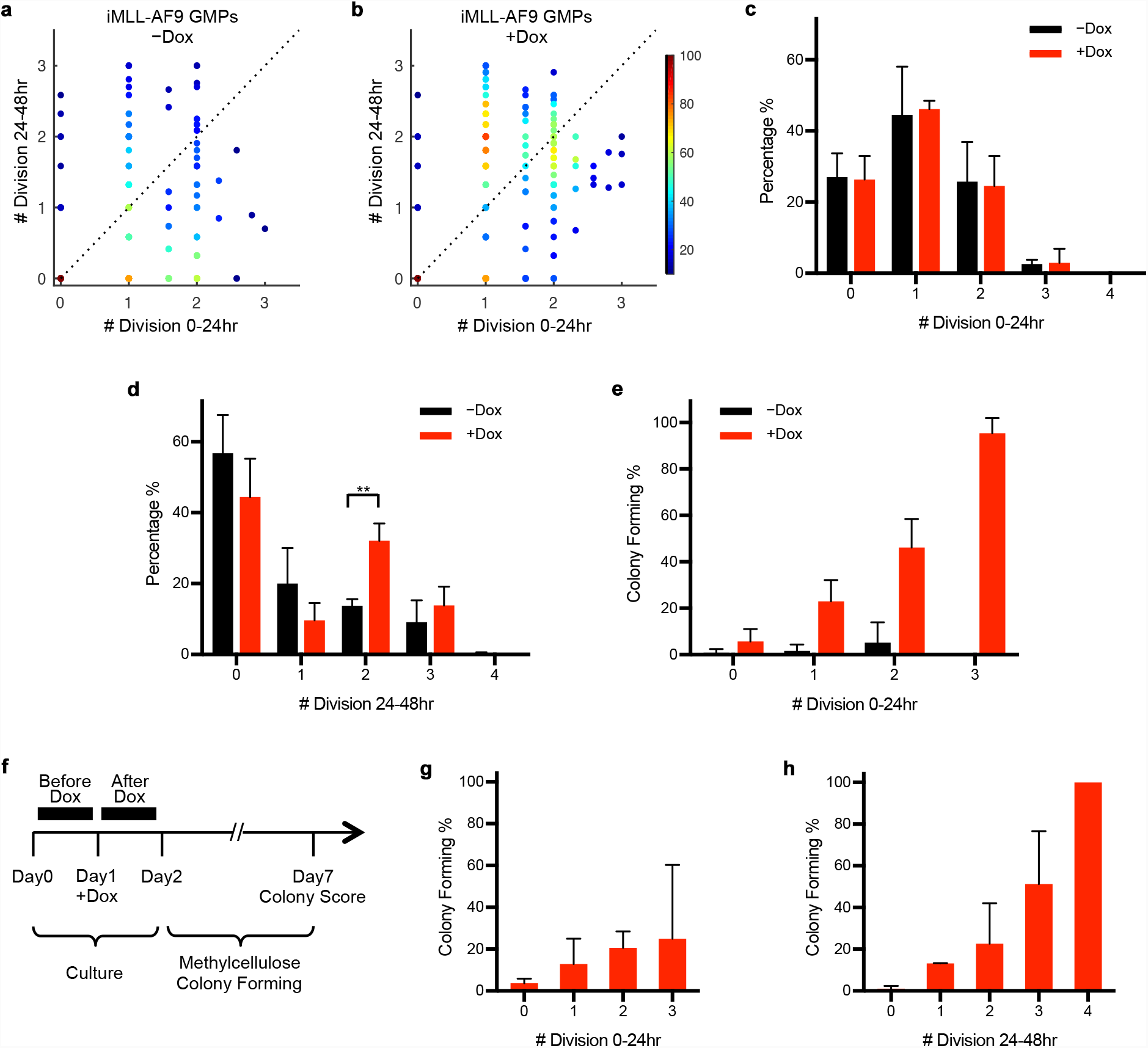
Permissiveness for MLL-AF9 mediated transformation is associated with the intrinsic cycling rate of GMPs. a, b, Cell cycle rates of iMLL-AF9 GMPs during the first 24 hours of culture (x axis) were plotted versus those of the second 24 hours (y axis). The presence of single GMPs was confirmed at O hour. The number of divisions during a 24-hour time widow was calculated as the Log2 value of cell number fold change. Each dot indicates an individual cell (a) n=274, -Dox; (b) n=378, +Dox (added at O hour). Pseudo color indicates normalized cell number distribution. Dotted diagonal line denotes y=x. c, Cell cycle rate distribution during the first 24 hours of iMLL-AF9 GMP culture. Cell cycle rates were determined as de scribed in a, b, and rounded up to the nearest integer. No more than 8 progeny were observed from a single GMP after the first 24 hours. The numbers of divisions likely represents an under-estimation due to potential co-occurring cell death. Detailed measurements are shown in Extended Data Table 1. d, Similar analysis as in c performed for the second 24 hours. e, Primary colony forming efficiency by iMLL-AF9 GMPs of different cycling rates, in the absence or presence of Dox. Cycling rates were defined by number of cell divisions during 0-24hr *in vitro* culture. f, Schematic timeline for measuring colony-forming efficiencies in relation to cell cycle rates, determined for the time before and after Dox addition at 24hr (Day1). g, Single cell colony forming efficiency in each subset of iMLL-AF9 GMPs of different cell cycle rates, which were deter mined by the number of divisions during 0-24hr of culture, prior to Dox induction, as shown in f. h, Single cell colony forming efficiency in each subset of iMLL-AF9 GMPs of different cell cycle rates, which were deter mined by the number of divisions during 24-48hr of culture, immediately after Dox induction, as shown inf.

To improve the throughput of this serial replating strategy which tracks malignancy initiation from single cells, we attempted to eliminate the pluck-and-replate steps. Specifically, we determined that culturing GMPs for two-days reduced the formation of non-malignant primary colonies to negligible levels (Fig. 1f, Extended Data Fig. 2d), likely due to differentiation during this time. The very few colonies that did form could not support serial re-plating (0 out of 30) (Fig. 1g). In contrast, many colonies emerged from the two-day cultured iMLL-AF9 GMPs when Dox was added (Fig. 1f), with the great majority of these supporting serial re-plating (86%, 74 out of 86) (Fig. 1g), and having up-regulated *Hoxa9* and *Meis1* (Fig. 1h), two well-established MLL-AF9 target genes^17,18^. These results demonstrate that the majority of the methylcellulose colonies developed from single iMLL-AF9 GMPs following a two-day culture were transformed.

This modified colony-forming assay enabled us to clonally track hundreds of individual GMPs, from their initial physiological states to when they first displayed *de novo* malignant phenotypes (Extended Data Fig. 2a, Fig. 1g-h). We determined that even though the GMPs initially appear identical and similarly express MLL-AF9 (Extended Data Fig. 1a), only ∼25% of them acquire malignancy (Fig. 1f). Importantly, this experimental system provided the opportunity to relate the normal cell behavior to their future fate outcome.

## Transformation initiates from cells in a naturally fast cycling state

We then asked whether the ∼25% GMPs acquiring malignancy were a random subset, or possessed specific cellular trait(s). We focused on their cell cycle rate, as cell cycle is a major contributor to cellular heterogeneity^19^ and an ultrafast cell cycle associates with cell fate plasticity^20^. We hypothesized that the transformation-permissive GMPs would display a distinct cell cycle rate. Alternatively, if malignancy arises randomly, the cell cycle rate of the transformation-permissive GMPs would resemble that of the bulk GMPs.

The cell cycle rate of single GMPs was determined by directly scoring the number of its progeny at 24 and 48 hours (Fig. 1d). After two days in culture, the cell cycle rate increased with Dox treatment (Fig. 2a,b). However, it is important to note that cell cycle rates remained identical within the first 24 hours between +/-Dox conditions (Fig. 2c, Extended Data Table 1), only diverging during the second day (Fig. 2d, Extended Data Table 1). Thus, the cell cycle rate during the first 24 hours reflects the intrinsic normal GMP behavior, while the second 24 hours includes the oncogene effects. The delay in cell cycle change within the first 24 hours of Dox treatment could be due to insufficient amount of MLL-AF9 being induced at this time, or the induced oncogene not significantly altering the cellular state early on.

With the ability to clonally track GMP cell cycle rates, we then asked whether the intrinsic cell cycle rate relates to its transformability. Indeed, the intrinsic cell cycle rate strongly correlated with their ability to form transformed colonies (Fig. 2e). Almost all GMPs that divided three times or more within the first 24 hours (representing ∼3% of total GMPs, or ∼0.006% of bone marrow nucleated cells) were transformed. The cell cycle rate of the second 24 hours of Dox treatment also correlated with transformation probability (Extended Data Fig. 2e), but this cell cycle rate no longer reflects the intrinsic ones and could be consequent to the prolonged oncogene activity (Fig. 2d). These data suggest that malignancy does not arise from random GMPs. Rather, the fastest cycling GMPs appear more probable to acquire malignancy.

The fast cycling behavior could underlie the high transformation probability itself, or simply mark the cell lineages with inherent ability to transform. If cell cycle rates mark distinct cell lineages, a fast cell cycle rate prior to Dox treatment should similarly identify transformation-competent GMPs. To test this, we delayed Dox addition for 24 hours (Fig. 2f) and assessed the cell cycle rates before (0-24 hours) and within the initial 24 hours of Dox treatment (24-48 hours) when cell cycle rates remain unaltered (Fig. 2c). Notably, the cell cycle rates prior to Dox treatment (0-24 hours) were minimally relevant for their transformation outcome (Fig. 2g). Instead, a much stronger correlation between the cell cycle rate at the time of Dox treatment (24-48 hours) and their transformation potential was detected (Fig. 2h). These data indicate that transformation permissiveness depends on the immediate proliferative state when MLL-AF9 is expressed. Because the proliferative history is less relevant for transformation permissiveness (Fig. 2g), these data also argue against the likelihood that additional replicative errors occurred during these rapid divisions are the major cause of their different propensities to transform.

## The early response to MLL-AF9 is consistent with the preservation of a GMP-like state

To understand how the immediate cellular state determines the permissiveness to transformation, we examined the earliest molecular response to MLL-AF9 induction (Extended Data Fig. 3a), when the cells still displayed their intrinsic proliferative behaviors. 24 hours of MLL-AF9 induction led to many differentially expressed genes (DEGs) between the +/–Dox GMPs (Fig. 3a). Most of the genes expressed higher in +Dox cells were involved in RNA metabolism related to protein translational processes, and most of the genes expressed lower in +Dox cells were immune function related (Fig. 3b). Surprisingly, the self-renewal program known to be reactivated by MLL-AF9^8^ was not detected at this time. Of the 41 L-GMP signature genes normally seen in HSCs, only 3 were expressed higher in our +Dox GMPs (Extended Data Fig. 3b). Furthermore, even though several key MLL-AF9 target genes (e.g. *HoxA9* and *Meis1*) were indeed higher with Dox treatment, their absolute expression levels were comparable to that of freshly-harvested GMPs and much lower than that in HSCs (Fig. 3c). Thus, reprogramming gene expression to a stem cell-like state was not the initial response to MLL-AF9. There were also down-regulated genes in +Dox GMPs as compared with – Dox GMPs, such as *Acod1* and *Cxcl2.* The expression of these genes was low in fresh GMPs, and Dox induction maintained their low expression level (Extended Data Fig.3c). Overall, these data suggest that MLL-AF9 dampens the gene expression changes accompanying the culture, and helps preserving the expression level of several genes to that of the originating GMPs.

**Fig 3:**
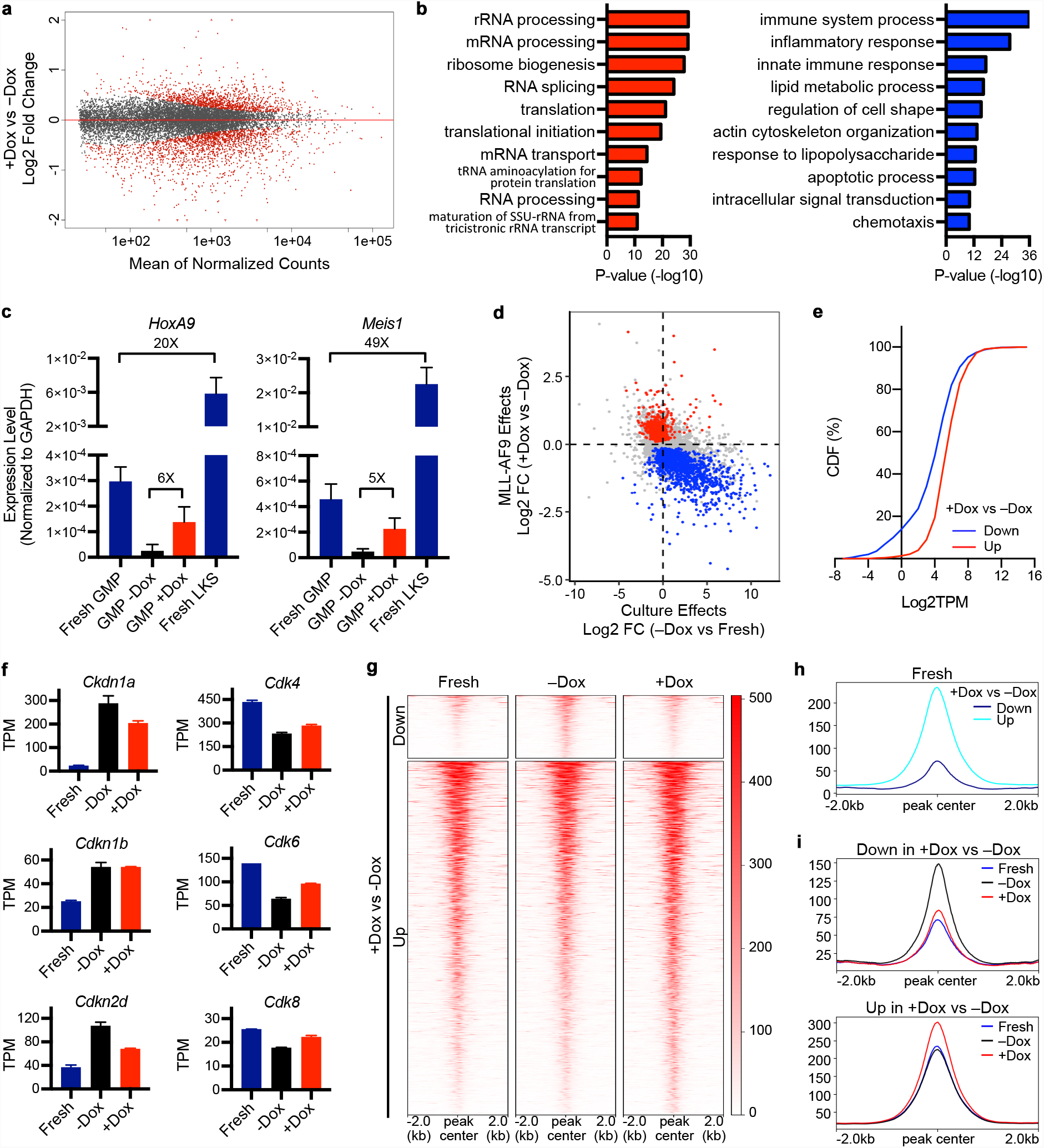
MLL-AF9 expression sustains the already-existing gene expression program in GMPs. a, MA plot showing gene expression changes in iMLL-AF9 GMPs treated +/-Dox for 24 hours. Red dots denote differen tially expressed genes (DEGs), p<0.05. b, Gene ontology (GO) analysis for DEGs as shown in a. The top 10 up-regulated (red bars) and down-regulated (blue bars) biological process terms in +Dox GMPs are shown. c, Comparison of *Hoxa9* and *Meis1* expression levels in samples as shown in a, plus freshly isolated LKS and GMPs. d, Scatter plot showing culture-induced gene expression changes (-Dox vs fresh GMPs) versus MLL-AF9 induced changes (+Dox vs -Dox GMPs). Blue dots represent the oncogene down-regulated DEGs; red dots represent the onco gene up-regulated DEGs. e, The DEGs shown in a display differential expression patterns in fresh GMPs. CDF plot of the expression levels of these DEGs in fresh GMPs is shown. Red line: genes up-regulated in +Dox samples; Blue line: genes down-regulated in +Dox samples. TPM: transcript per million. f, Representative expression levels of up-/down-regulated cell cycle genes. g, Heatmap showing ATAC-seq peak intensities in iMLL-AF9 GMPs, fresh or +/-Dox for 24 hours. Regions Down (697) or Up (3929) were defined by comparing +Dox GMPs with -Dox GMPs. h, The Up and Down ATAC-seq peaks, as defined in g, display differential accessibility in fresh GMPs. Meta plot sum marizing their intensities in fresh GMPs is shown. i, Dox inductions (red line) prevented the opening of chromatin regions during GMP culture (upper, blue vs black lines), and promoted the further opening of chromatin regions that are already accessible in fresh GMPs (lower, blue line). The ATAC-seq peaks (Down or Up) are as defined in g.

To determine whether MLL-AF9 dampens the gene expression changes in general, we plotted our Dox-induced gene expression changes against the gene expression changes elicited by culture (i.e.–Dox GMPs vs fresh GMPs) (Fig. 3d). This revealed that MLL-AF9 induction antagonized culture-induced gene expression changes overall. On absolute gene expression levels, Dox up-regulated DEGs expressed at higher levels in fresh GMPs, while Dox down-regulated DEGs expressed lower (Fig.3e). These results indicate that the preservation of gene expression by MLL-AF9 is widespread, and the expression state of a given gene determines its initial response to MLL-AF9. Specifically, an actively expressed gene is likely to retain its high expression while one with lower expression continues to remain low. Taken together, the early gene expression changes in response to MLL-AF9 were consistent with the preservation of their expression status in fresh GMPs. Of note, expression of cell cycle genes, including CDK6, which is essential for transformation by this oncogene^21^, was preserved at levels similar to fresh GMPs (Fig. 3f). The effects on cell cycle gene expression may be related to how fast cell cycle rate contributes to MLL-AF9 mediated transformation (Fig. 2e).

In agreement with the gene expression data, increased ATAC-seq signals were detected with up-regulated genes, such as *HoxA9* (Extended Data Fig. 3d), and vice versa with decreased ATAC-seq signals in down-regulated genes, such as *Acod1* (Extended Data Fig. 3e). The differentially accessible regions produced many similar GO terms (Extended Data Fig. 3f) as those of the DEGs (Fig. 3b). Importantly, the genomic regions corresponding to the Dox-increased ATAC-seq peaks were already accessible in normal/fresh GMPs, while the regions corresponding to Dox-decreased ATAC-seq peaks had low accessibility in normal GMPs (Fig. 3g,h). With fresh GMPs as a reference, MLL-AF9 expression appears to have prevented culture-induced chromatin region opening (Fig. 3i, upper panel), and enhanced the accessibility of those already open regions (Fig. 3i, lower panel), particularly so around its direct target sites^22^ (Extended Data Fig. 3g,h). Overall, MLL-AF9 expression in GMPs upholds the GMP state at the chromatin level.

Taken together, the earliest gene expression response to MLL-AF9 recapitulates a cell state resembling the naturally existing GMPs. Viewed in this light, MLL-AF9 could transform cells by preserving and perpetuating a rapidly proliferating immature myeloid cell program, one that is inherent to a subset of normal GMPs.

## MLL-AF9 helps to preserve the gene expression programs in additional cell types

To determine whether MLL-AF9 helps to sustain gene expression programs in general, we induced its expression in additional cell types. This is relevant because expressing MLL-AF9 in hematopoietic stem and progenitors upstream of GMPs give rise to a similar disease *in vivo*^7^, which could occur by two potential means. (1) MLL- AF9 reprograms gene expression, yielding the same target malignancy irrespective of the initial cell type, in a manner similar to pluripotency transcription factor induced reprogramming^23^; or (2) MLL-AF9 facilitates the continuation of a proliferative progenitor state that is similar to the transformed state.

To determine whether MLL-AF9 induces common target genes across different cells, we treated hematopoietic stem and progenitor cells (Lin-c-Kit+ Sca-1+, LKS cells), GMPs, and differentiated myeloid cells (Mac1+) with Dox for 2 days (Fig. 4a). While MLL-AF9 was similarly induced in all cells (Extended Data Fig. 4a), Dox led to distinct proliferative responses (Fig. 4b, Extended Data Fig. 4b). GMPs, which are naturally proliferative^24^, increased their proliferation in the presence of Dox. In contrast, neither LKS nor Mac1+ cells did so, coinciding with their slower or absence of proliferation. Accordingly, only the Dox-treated GMPs, but not the LKS or Mac1+ cells, displayed consistent positive enrichment in proliferation gene sets (Fig. 4c). Furthermore, although Dox failed to enhance the proliferation of freshly isolated LKS cells (Extended Data Fig. 4c), it did so when LKS were first activated into cycling from a largely quiescent state (Extended Data Fig. 4d)^20^. The differential responses in fresh and cultured LKS cells further support the importance of initial cellular states in determining MLL-AF9 effects. These data are consistent with a previous report for MLL-ENL expressing HSCs^13^, suggesting functional conservation among MLL-fusion oncogenes. Thus, MLL-AF9 expression does not impart a universal proliferative response (Fig. 4b-c, Extended Data Fig. 4b-d). Rather, the effects depend on the specific proliferative state of the starting cells.

**Fig 4:**
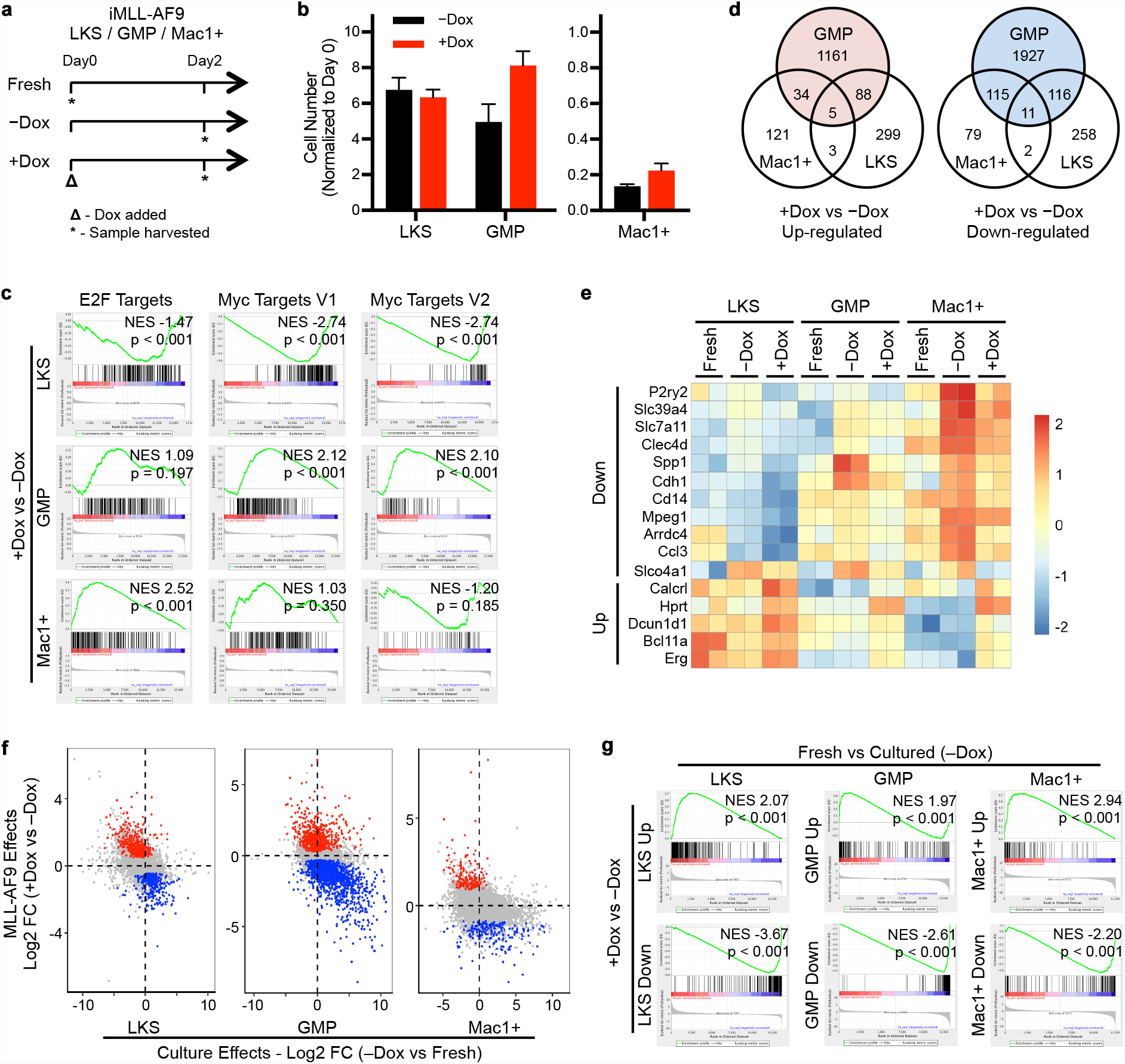
The primarygene expression changes in response to MLL-AF9 induction is cell-context dependent. a, Workflow showing iMLL-AF9 LKS, GMP, or Mac1+ cells after their isolation on Day 0 (Fresh), or treated +/-Dox for two days. b, Proliferation rates of respective cell types during 2-days of culture +/-Dox. c, Distinct proliferative responses to MLL-AF9 induction in LKS, GMP and Mac1+ cells, as determined by Gene Set En richment Analysis (GSEA). Note GMPs displayed consistent positive enrichment for all cell cycle gene sets. In contrast, the response from LKS was all negative and the response from Mac1+ cells was not consistent. d, Venn diagram showing little overlap among up-/down-regulated DEGs from LKS, GMP and Mac1+ cells. Cell type-specific DEGs (p<0.05) were defined between the respective cells treated +/-Dox. e, Heatmap showing expression levels of 11 commonly down-regulated and 5 commonly up-regulated genes in all three cell types. Log2TPM of each gene was row normalized. f, Scatter plot showing culture-induced gene expression changes (-Dox vs fresh) versus MLL-AF9 induced changes (+Dox vs -Dox) in three cell types. Blue dots represent the oncogene down-regulated DEGs; red dots represent the on cogene up-regulated DEGs g, The Dox-induced DEGs, similar to those defined ind but at p<0.01, were queried against the gene expression chang es occurred during culture in the respective cell types by GSEA. For GMPs, only the top 200 DEGs were used.

In agreement with the disparate proliferative responses, there was minimal overlap amongst the Dox-induced DEGs across the three cell types, with only 5 genes being commonly up-regulated and 11 commonly down-regulated (Fig. 4d,e). One of the commonly up-regulated genes is *Hprt*, the host locus for the iMLL-AF9 allele (Fig. 1a), corroborating successful transgene induction in all cell types from this locus (Extended Data Fig. 4a). Taken together, the primary MLL-AF9-responsive genes differ according to the specific cell types in which it is expressed. An early universal gene signature induced by MLL-AF9 was not present.

Contrasting the lack of common target genes across different cell types, MLL- AF9 expression invariably countered the culture-elicited gene expression changes (Fig. 4f,g). In all three cell types, MLL-AF9 up-regulated genes were significantly enriched in the respective fresh cells, while MLL-AF9 down-regulated ones were negatively enriched (Fig. 4f,g). The enrichment was specific to the respective cell type itself and no enrichment across cell types was detected (Extended Data Fig. 4e). Thus, although MLL-AF9 failed to induce a common gene signature, it did sustain the existing gene expression programs in multiple cell types.

## Modification of the initial cell state by transient cell cycle inhibition mitigates MLL-AF9 mediated transformation

The above model implies that modifying the initiating cell cycle state, even by transient inhibition, could alter MLL-AF9’s oncogenic effects on gene expression, and result in lasting reduction in transformation. To test this possibility, we treated GMPs with palbociclib (PD0332991), a CDK4/6 inhibitor^25^ (Fig. 5a). At 500nM, palbociclib mildly and reversibly decreased GMP proliferation (Fig. 5b, Extended Data Fig. 5a-c). It indeed decreased expression of many cell cycle related genes (Fig. 5c,d, Extended Data Fig. 5d), without significant changes in apoptosis-associated genes (Extended Data Fig. 5e). We then analyzed the palbociclib-modified gene expression changes in response to MLL-AF9 (Fig. 5a), and found that palbociclib dampened the response to MLL-AF9 expression, resulting in fewer Dox-induced DEGs (Fig. 5e, Extended Data Fig. 5f). In the presence of palbociclib, MLL-AF9 failed to up-regulate a significant subset of proliferation-related genes (Fig. 5f), and lost suppression of many differentiation-related genes (Fig. 5g). Furthermore, many of the changes that did pass as Dox-induced DEGs were subdued in the presence of palbociclib (Fig. 5h). Taken together, these data suggest that even mild cell cycle inhibition could dampen MLL-AF9’s effect, as assessed by gene expression changes.

**Fig 5:**
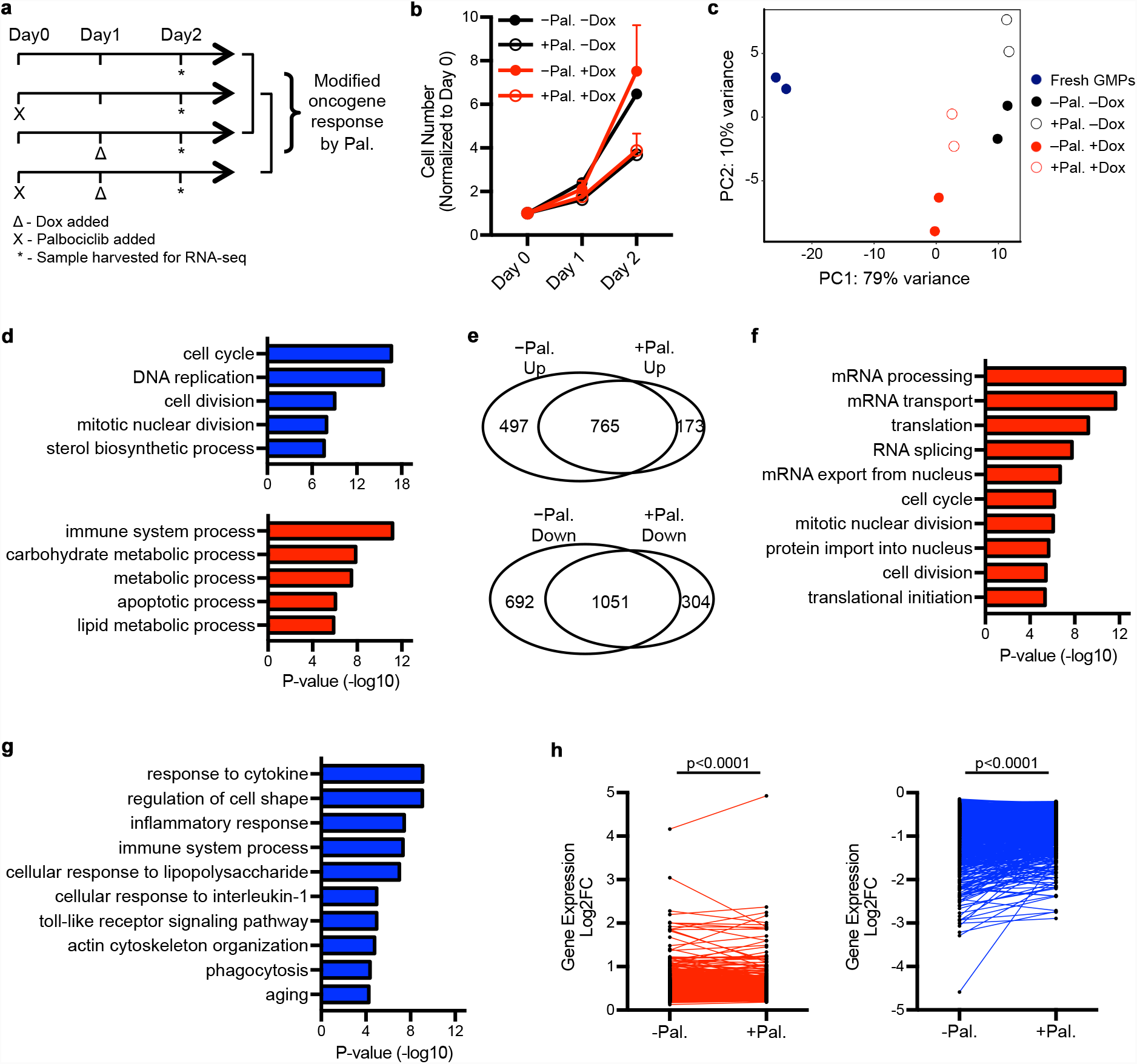
Modification of the initial cell state by cell cycle inhibition mitigates MLL-AF9 mediated changes in gene expression. a, Timeline schematics illustrating treatment with a CDK4/6 inhibitor palbociclib or vehicle control (+/-Pal.) for iMLL-AF9 GMPs, prior to their culture, in the presence or absence of Dox (+/-Dox). All cells were harvested for RNA-seq analyses after a total of 2 days culture. Comparisons used for analyses were indicated by the brackets on the right. Fresh GMPs were also included in the analysis. b, The proliferation rates of iMLL-AF9 GMP samples used for RNA-seq., confirming palbociclib effects on proliferation after two days. c, Principal Component Analysis (PCA) showing Dox-induced gene expression changes, as modified by Palbociclib treatment. d, Top 5 GO biological process terms of DEGs (p<0.01) between iMLL-AF9 GMPs treated with palbociclib or vehicle control (+/-Pal.). No Dox was added in these samples. Red and blue bars denote genes up- or down-regulated in +Pal. GMPs, respectively. e, Venn diagrams showing the Dox-induced DEGs when palbociclib was present (+Pal.) or not (-Pal.). f, GO analysis for Dox up-regulated DEGs only in the absence of palbociclib (497 genes as shown in e). The top 10 bio logical process terms are shown. g, GO analysis for Dox down-regulated DEGs only in the absence of palbociclib (692 genes as shown in e). The Top 10 biological process terms are shown. h, Dox-induced gene expression changes (Log2 fold change) in the absence or presence of palbociclib. The 765 com monly up-regulated DEGs in e are shown in red, and the 1051 commonly down-regulated DEGs in e are shown in blue. Each dot denotes an individual gene and the same genes in +/-Pal. conditions are connected by a line. P value was cal culated by paired t test.

We next determined whether this mild cell cycle inhibition reduces permissiveness to transformation (Fig. 6a). Palbociclib treatment preferentially abrogated the most proliferative GMP subsets, with the cells that could divide three times or more within 24 hours becoming barely detectible (Fig. 6b, Extended Data Table 2). Tracking the cells undergoing palbociclib treatment in micro-wells revealed that their ability to form transformed colonies was decreased (Fig. 6c, Extended Data Fig. 6a). Palbociclib treatment of iMLL-AF9 GMPs also significantly decreased transformed colony formation (by ∼50%) in bulk methylcellulose cultures, where colony size is not restricted by the micro-well (Fig. 6d, Extended Data Fig. 6b-c). Importantly, the size of the persisting transformed colonies did not decrease (Fig. 6e-f), suggesting palbociclib reduced the units of transformation, but not the general proliferative capacity of the still transformation competent cells.

**Fig 6:**
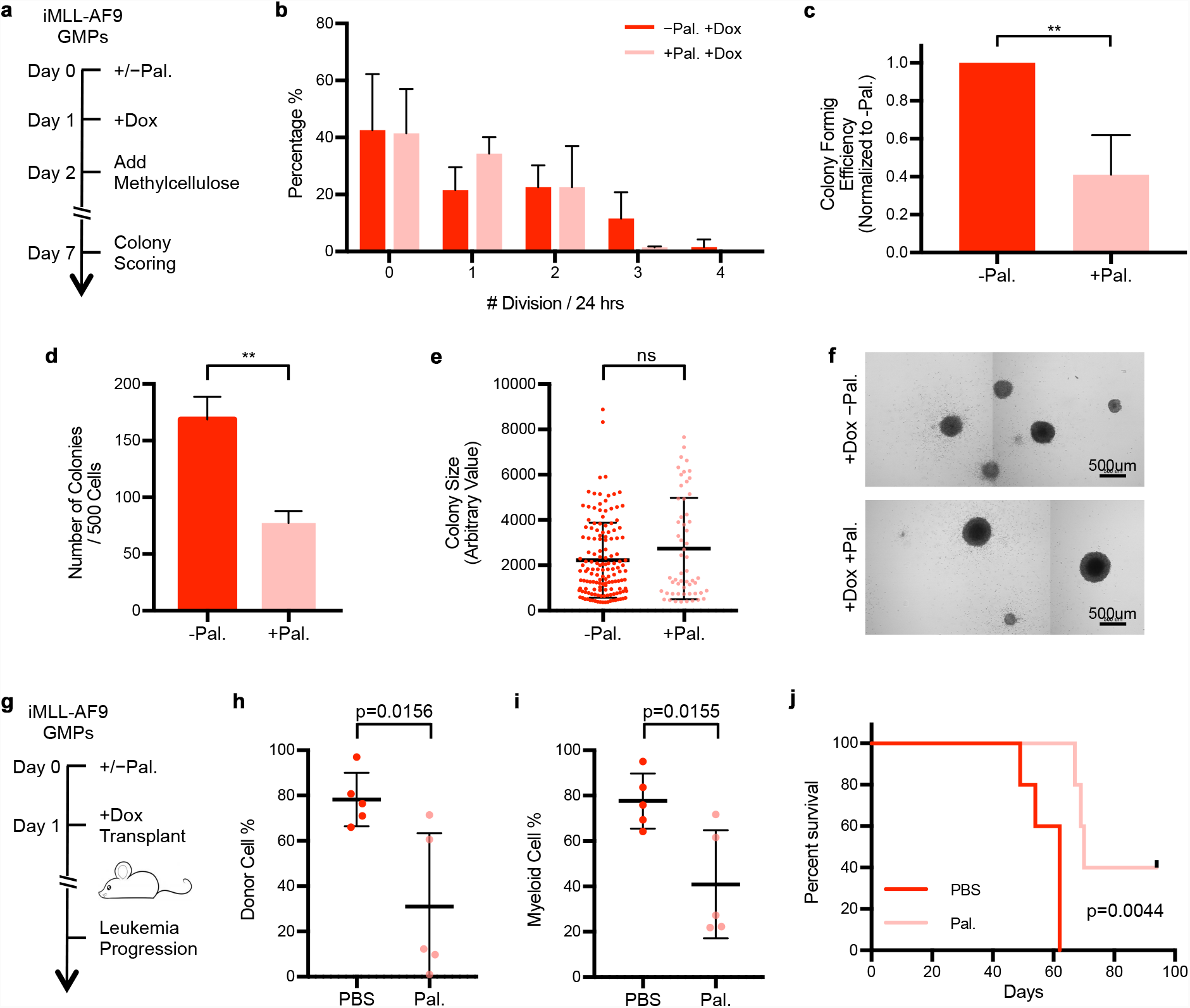
Modification of the initial cell state by mild cell cycle inhibition mitigates MLL-AF9 mediated transformation. a, Schematics illustrating the formation of transformed colonies by iMLL-AF9 GMPs in the presence or absence of pal bociclib (+/-Pal.). b, Cell cycle rate distribution of iMLL-AF9 GMPs cultured in micro-wells in the presence or absence of palbociclib (+/-Pal.). Palbociclib was added at Ohr. Dox was added at 24hr, and cell cycle rate was determined from 24hr to 48hr. Error bars represent standard deviation. Data is pooled from 4 (-Pal.) and 3 (+Pal.) independent experiments. Detailed measurements are shown in Extended Data Table 2. c, Quantification of transformed colony forming efficiencies in micro-wells by control iMLL-AF9 GMPs (-Pal), or when Palbociclib was added from Ohr. For each experiment, the portion of Pal-treated iMLL-AF9 GMPs forming transformed colonies were normalized to control iMLL-AF9 GMPs. Dox was added to all samples at 24hr. Data is pooled from three independent experiments. d, e, Quantification of colony number (d) or size (e) of bulk cultured iMLL-AF9 GMPs in methylcellulose in the absence and presence of palbociclib. Dox was added to all conditions. To prevent proliferation rate recovery after palbociclib washout (Extended Data Fig. 5c), we maintained palbociclib at 250nM after treating the cells at 500nM for the first 2 days (Extended Data Fig. 6c,d) f, Representative images showing colony morphologies from iMLL-AF9 GMPs +/-Pal.. Scale bar: 500um. g, Schematics illustrating *in vivo* leukemogenesis by iMLL-AF9 GMPs. iMLL-AF9 GMPs were treated with palbociclib *in vitro* for one day (+/-Pal.) prior to their injection into sub-lethally irradiated recipient mice, which received three injec tions of palbociclib or vehicle control (+/-Pal.) on day 0, day 1 and day 2. All mice received 20,000 GMPs and were fed with Dox water. h, Percentage of donor-derived cells (CD45.1 /2) in the peripheral blood of recipient mice (CD45.2), assayed 7 weeks post transplantation. i, Percentage of myeloid cells (Mac1+) in the peripheral blood of recipient mice, assayed 7 weeks post transplantation. j, Kaplan-Meier curve of recipient mice developing AML after iMLL-AF9 GMP transplantations.

To examine whether transient palbociclib treatment leads to lasting reduction in transformation *in vivo*, we transplanted iMLL-AF9 GMPs at a dosage where all mice develop AML when fed with Dox (Fig. 6g). Cohorts of Dox-fed mice were briefly treated with either vehicle control or palbociclib at a previously validated dosage^26^, immediately following iMLL-AF9 GMP injection (Fig. 6g, Extended Data Fig. 6d). Similar to the results *in vitro*, this mild, transient palbociclib treatment reduced transformation *in vivo*, leading to decreased leukemia cell burden and prolonged animal survival (Fig. 6h-j). A subset of the palbociclib-treated mice never developed AML and remained healthy for the entire time studied (Fig. 6j). Together, these results demonstrate that transient modification of the cellular state by cell cycle inhibition could effectively reduce the likelihood of malignancy emergence *in vivo*.

## Discussion

Because malignant transformation is usually a protracted and rare event, it has been difficult to experimentally determine which cells among those bearing cancer predisposing mutation(s) acquire malignancy. We overcame this limitation by establishing an inducible MLL-AF9 allele combined with a system to track transformed fate outcome at clonal level. We identified a cellular state in which the presence of this single oncogene makes transformation nearly a certainty. At least in this oncogenic model, the proliferative state is not the consequence of malignant transformation, but rather a prerequisite for its initiation. Based on the observation that MLL-AF9 helps to preserve the ongoing gene expression program across multiple cellular contexts, we propose that the rapidly proliferating, poorly differentiated myeloid progenitor cell state undergoes transformation when it is perpetuated by MLL-AF9 expression, fulfilling the functional definition of malignancy. Our data are in agreement with the rapidly cycling myeloid progenitors to be the immediate cell-of-origin^13,27–29^, and suggest HSCs to be more probable in serving as the reservoir to sustain the rapidly cycling progenitor compartment. The heightened permissiveness to transformation by faster cycling cells may underlie the myeloproliferative phase preceding many AMLs, possibly by increasing the number of cells permissive of transformation.

Our model depicting MLL-AF9 sustaining ongoing gene expression programs of fast cycling cells implies that mitotic bookmarking mechanisms^30,31^ might be particularly relevant for MLL-fusion oncogenes. MLL can bind to mitotic chromatin marking the regions for rapid transcriptional reactivation in the next cell cycle, although it also binds to interphase chromatin^32^. Future studies should address whether MLL-AF9’s effectiveness in fast cycling cells also stems from the oncoprotein’s preference for mitotic chromatin. Nonetheless, our model could easily explain why MLL-fusion leukemias are sensitive to BET inhibitors^33,34^ and CDK4/6 inhibitors^21,35^, as Brd4 is a component of the mitotic bookmarking machinery^36^ and CDK4/6 inhibition halts G1/S progression^37^, respectively. Besides malignancy, the same rapidly proliferating GMPs are also extraordinarily efficient in initiating pluripotency^20^, although the mechanisms responsible for these two processes should be distinct. Acquisition of pluripotency represents a departure from the somatic state while MLL-fusion oncogene mediated transformation perpetuates it. This might be the underlying reason why the same Dot1L inhibitor promotes somatic cell reprogramming^38^ but inhibits MLL-fusion leukemia^39,40^.

## Materials and Methods

### Mice

All mouse work has been approved by the Institutional Animal Care and Use Committee (IACUC) of Yale University. All mice used in this study were maintained at Yale Animal Resources Center (YARC). To generate the inducible MLL-AF9 knock-in mouse, the cDNA encoding human MLL-AF9 linked with IRES-NGFR^14^ was targeted into the A2Lox.cre mESC cell line as previously described^15^. Correctly targeted mESCs were injected into E3.5 blastocysts by Yale Genome Editing Center. The knock-in mice were crossed with a Rosa26 rtTA allele^16^, and bred to reach homozygosity for both MLL-AF9 and rtTA alleles. Mouse genotyping was determined by PCR using primers listed in Extended Data Table 3.

### FACS Sorting and Analysis

Antibodies used in this study are listed in Extended Data Table 4. GMPs were isolated using fluorescence activated cell sorting (FACS) on BD Aria as defined previously (Lin-cKit+Sca-1-CD34+CD16/32+)^20,41^. Hematopoietic stem and progenitor cells upstream of GMP were sorted as LKS (Lin-cKit+Sca-1+), and differentiated myeloid cells were sorted as Mac1+ from Lin+ populations. FACS analyses were done using BD LSRII and analyzed with FlowJo.

For Hochest DNA content staining, cells were fixed in 70% ethanol for 30 minutes on ice, followed by staining with 5ug/mL Hochest at room temperature for 15 minutes, and analyzed on LSRII directly.

### Cell Culture

Freshly isolated GMPs were cultured in x-vivo15 (Lonza, 04-418Q), supplemented with 50ng/ml mIL3, 50ng/ml Flt3L, 50ng/ml mTPO, and 100ng/ml mSCF. 2ug/ml Dox (Sigma, D9891) was used for MLL-AF9 oncogene induction. Palbociclib (Selleckchem, S1579) dissolved in H_2_O was added to culture medium, and used at concentrations as indicated.

### Methylcellulose Colony Forming Assay

GMPs were plated in MethoCult GF M3434 (Stem Cell Technologies) for colony forming, following manufacture’s instructions. For serial colony forming, colonies were scored after 7 days from initial plating, cells were collected, and re-plated as single cell suspension.

For tracking colony formation at clonal level, micro-wells were casted using 1.2% low gelling temperature argarose (Sigma, A9045) with MicroTissues^®^ 3D Petri Dish^®^ (Sigma, Z764043). The micro-well gels were submerged in PBS for equilibration at 37°C for twenty minutes, followed by x-vivo15 overnight. Medium was removed the next day, and GMPs resuspended at 2,500 cells/mL in 75ul were loaded into each gel chamber, which were then incubated at 37°C for fifteen minutes to allow cells settling down into individual wells. Afterwards, extra medium supplemented with cytokines was added to cover the entire gels in order to provide sufficient volume and nutrients for cell growth. After two days in liquid culture, medium was replaced with MethoCult GF M3434 for colony growing. Schema is shown in Extended Data Fig. 2a.

### Single GMP Cell Cycle Rate Density Plot

At each data point, a circle with radius 0.5 is used to determine area and data points included to calculate the local density. The color scale represents data density. The color axes of Fig. 2a and Fig. 2b are scaled to the same range.

### Image Acquisition and Processing

For Giemsa images (Extended Data Fig.1d), peripheral blood and bone marrow smears were stained with May-Grunwald Giemsa (Sigma MG500), and images were taken in bright field under 60x objective, using Olympus BX51TRF.

Colonies and micro-wells images were acquired using ImageXpress Micro 4 high-content imaging system (Molecular Devices) at 4x or 10x objective. Images from adjacent fields were then stitched together in ImageJ to generate the whole micro-well views. Colony size was quantified using MetaMorph image analysis software.

### GMP Transplantation and Induction of AML ***in vivo***

FACS sorted GMPs were transplanted through tail vein into recipient mice that had received 6-Gy irradiation with gamma source. If oncogene was to be induced, 0.1 mg of doxycycline was given via i.p. injection at the time of transplantation. Animals were then fed with 1g/L doxycycline in drinking water sweetened with 10g/L sucrose. Development of leukemia was monitored by periodic analysis of peripheral blood obtained by tail vein bleeding.

### Palbociclib Treatment ***in vivo***

For *in vivo* palbociclib administration, palbociclib was dissolved in PBS and given by i.p. injection at 30 mg/kg. Recipient mice were given three shots of palbociclib, one-day prior to, the day of, and one-day after the transplantation. Schema is shown in Extended Data Fig. 6d.

### RNA Extraction, Reverse Transcription and QPCR

Total RNA was extracted with TRIzol^®^ Reagent (Ambion) and reverse transcribed using SuperScript™ III First-Strand Synthesis SuperMix (Invitrogen) following manufacturer’s instructions. Quantitative real-time PCR was performed using the iQ™ SYBR^®^ Green Supermix (Bio-Rad). Gene expression levels were normalized to GAPDH level in the same sample. Gene specific primers are listed in Extended Data Table 3.

### RNA-Seq and Data Analysis

The quality of total RNA was analyzed on Agilent Bioanalyzer. The RNA samples with more than 8 RNA intergration number (RIN) were chosen for RNA-Seq library preparation using TruSeq Stranded mRNA Library Preparation Kit from Illumina (Cat # RS-122-2101). GMP samples treated with Dox for 24 hours (Fig. 3, 5) were sequenced on HiSeq 2000 platform, and LKS/GMP/Mac1+ samples treated with Dox for two days (Fig. 4) were sequenced on HiSeq 4000 platform as pair-end 100 cycles following manufacture’s instruction. Sequencing reads were mapped to mouse mm10 using TopHat. Gene counts were quantified by Featurecounts and differentially expressed genes (DEGs) were identified using DeSeq2, with adjusted P value < 0.05. Gene ontology (GO) analysis of DEGs was performed through DAVID (https://david.ncifcrf.gov/). Scatter plots and heatmap were generated using ggplot2. Gene Set Enrichment Analysis (GSEA) was performed using GSEA software with default settings.

### ATAC-Seq and Data Analysis

ATAC libraries were prepared as previously described^42^. 50,000 cells were used per reaction for each cell type. Libraries were sequenced on Illumina HiSeq 2500 platform at Yale Center for Genome Analysis (YCGA). Sequencing reads were trimmed for adapter sequences using Cutadapt and aligned to mouse mm10 using Bowtie2. Duplicates were removed using Picard from uniquely mapped reads, and MACS2 BAMPE mode was used to call peaks. Differential peaks were analyzed using Diffbind default settings, and visualizations were done using deepTools (version 2.5.7). GO analysis was performed using GREAT (http://great.stanford.edu/public/html/index.php).

### Statistical Analysis

Student t-test was used in all the statistical analysis. Log-rank (Mantel-Cox) test was used on Kaplan-Meier survival curves.

## Data Availability

The Gene Expression Omnibus (GEO) accession number for RNA-Seq and ATAC-Seq is GSE121768.

## Acknowledgements

This work was supported by Gilead Sciences; Charles Hood Foundation and NIH/DP2.

---

**Extended Data Table 4.** Flow cytometry antibodies

| **Antibody** | **Clone** | **Vendor** |
| --- | --- | --- |
| CD271 (NGFR) - APC | ME20.4 | BioLegend® |
| Ly-6G/6C - Biotin | RB6-8C5 | BD Pharmingen™ |
| CD3e - Biotin | 145-2C11 | BD Pharmingen™ |
| CD45R/B220 - Biotin | RA3-6B2 | BD Pharmingen™ |
| CD11b - Biotin | M1/70 | BD Pharmingen™ |
| Ter119 - Biotin | Cat. 553672 | BD Pharmingen™ |
| CD8a - Biotin | 53-67 | BD Pharmingen™ |
| CD4 - Biotin | GK1.5 | BD Pharmingen™ |
| CD117 - APC | 2B8 | BD Pharmingen™ |
| Ly-6A/E (Sca-1) - PE | D7 | eBioscience |
| Streptavidin – BV510 | Cat. 563261 | BD Horizon™ |
| CD16/32 – PE-Cy7 | 93 | eBioscience |
| CD34 – Alexa Fluor® 700 | RAM34 | BD Pharmingen™ |
| CD11b - APC | M1/70 | eBioscience |
| CD45.1 – BV711 | A20 | BD Horizon™ |
| CD45.2 – Pacific Blue™ | 104 | BioLegend® |

**Extended Data Table 3.**
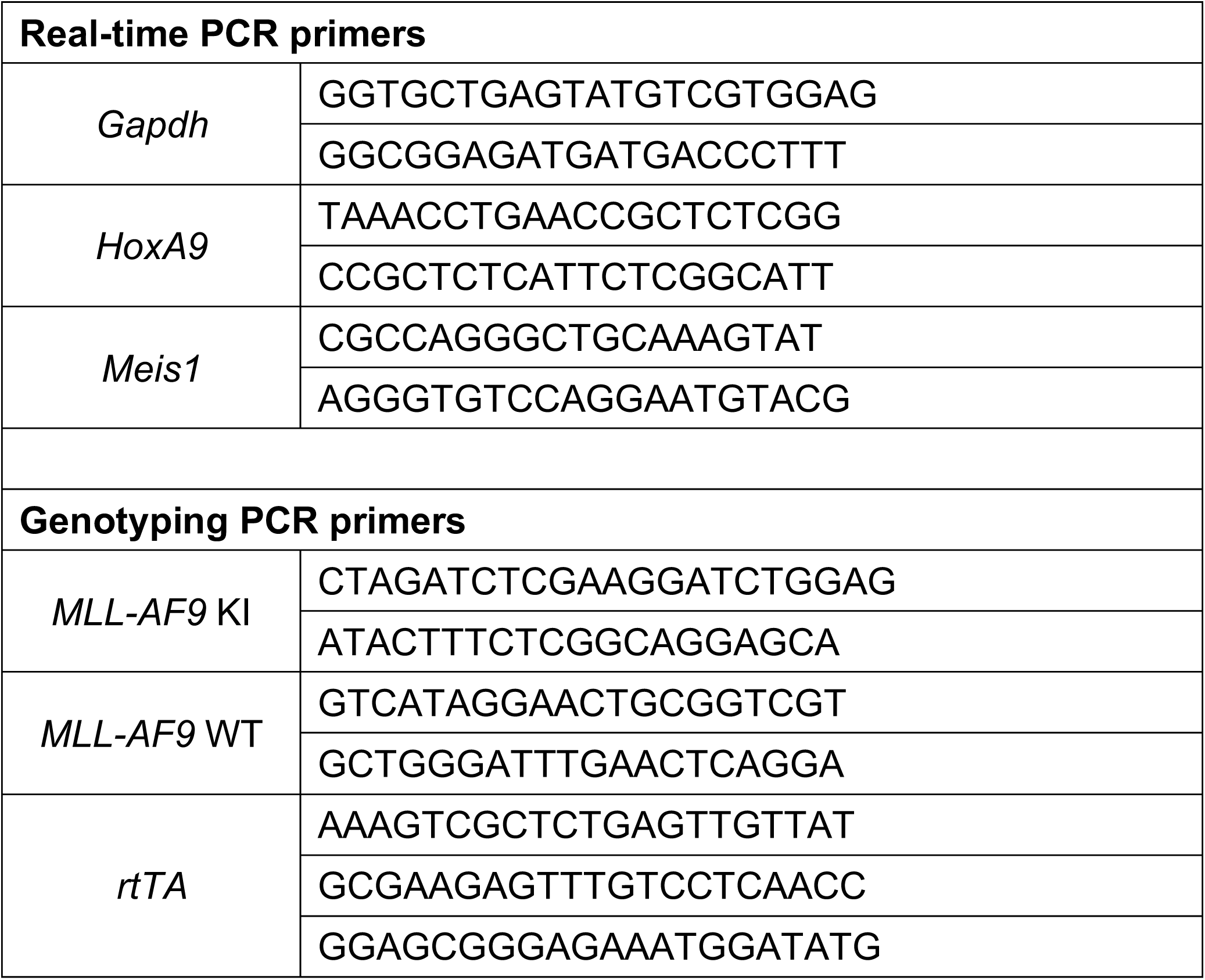
Oligo sequences

**Extended Data Table 2.** Single-cell cell cycle measurements (+/-Pal.)

| **-Pal.** | **Total cells** | **# Division / 24hr** |  |  |  |  |
| --- | --- | --- | --- | --- | --- | --- |
|  |  | 0 | 1 | 2 | 3 | 4 |
| *Rep. 1* | 87 | 55<br>(63.2%) | 15<br>(17.2%) | 11<br>(12.6%) | 6<br>(6.9%) | - |
| *Rep. 2* | 137 | 44<br>(32.1%) | 46<br>(33.6%) | 33<br>(24.1%) | 13<br>(9.5%) | 1<br>(0.7%) |
| *Rep. 3* | 179 | 37<br>(20.7%) | 31<br>(17.3%) | 56<br>(31.3%) | 45<br>(25.1%) | 10<br>(5.6%) |
| *Rep. 4* | 125 | 68<br>(54.4%) | 23<br>(18.4%) | 28<br>(22.4%) | 6<br>(4.8%) | - |

| **+Pal.** | **Total cells** | **# Division / 24hr** |  |  |  |  |
| --- | --- | --- | --- | --- | --- | --- |
|  |  | 0 | 1 | 2 | 3 | 4 |
| *Rep. 1* | 151 | 85<br>(56.3%) | 44<br>(29.1%) | 20<br>(13.2%) | 2<br>(1.3%) | - |
| *Rep. 2* | 158 | 40<br>(25.3%) | 53<br>(33.6%) | 62<br>(39.2%) | 3<br>(1.9%) | - |
| *Rep. 3* | 79 | 34<br>(43.0%) | 32<br>(40.5%) | 12<br>(15.2%) | 1<br>(1.3%) | - |

**Extended Data Table 1.** Single-cell cell cycle measurements (+/-Dox)

| **-Dox** | **Total cells** | **# Division 0-24hr** |  |  |  |  |
| --- | --- | --- | --- | --- | --- | --- |
|  |  | 0 | 1 | 2 | 3 | 4 |
| *Rep. 1* | 75 | 15<br>(20%) | 44<br>(58.7%) | 13<br>(17.3%) | 3<br>(4%) | - |
| *Rep. 2* | 60 | 20<br>(33.3%) | 26<br>(43.3%) | 13<br>(21.7%) | 1<br>(1.7%) | - |
| *Rep. 3* | 133 | 37<br>(27.8%) | 42<br>(31.6%) | 51<br>(38.3%) | 3<br>(2.3%) | - |

| **+Dox** | **Total cells** | **# Division 0-24hr** |  |  |  |  |
| --- | --- | --- | --- | --- | --- | --- |
|  |  | 0 | 1 | 2 | 3 | 4 |
| *Rep. 1* | 83 | 28<br>(33.7%) | 40<br>(48.2%) | 15<br>(18.1%) | - | - |
| *Rep. 2* | 148 | 36<br>(24.3%) | 69<br>(46.6%) | 32<br>(21.6%) | 11<br>(7.4%) | - |
| *Rep. 3* | 147 | 31<br>(21.1%) | 64<br>(43.5%) | 50<br>(30.4%) | 2<br>(1.4%) | - |

| **-Dox** | **Total cells** | **# Division 24-48hr** |  |  |  |  |
| --- | --- | --- | --- | --- | --- | --- |
|  |  | 0 | 1 | 2 | 3 | 4 |
| *Rep. 1* | 75 | 47<br>(62.7%) | 15<br>(20%) | 11<br>(14.7%) | 2<br>(2.7%) | - |
| *Rep. 2* | 60 | 38<br>(63.3%) | 6<br>(10%) | 7<br>(11.7%) | 9<br>(15%) | - |
| *Rep. 3* | 133 | 59<br>(44.4%) | 40<br>(30.1%) | 20<br>(15.0%) | 13<br>(9.8%) | 1<br>(0.8%) |

| **+Dox** | **Total cells** | **# Division 24-48hr** |  |  |  |  |
| --- | --- | --- | --- | --- | --- | --- |
|  |  | 0 | 1 | 2 | 3 | 4 |
| *Rep. 1* | 83 | 47<br>(56.6%) | 7<br>(8.4%) | 22<br>(26.5%) | 7<br>(8.4%) | - |
| *Rep. 2* | 148 | 60<br>(40.5%) | 8<br>(5.4%) | 52<br>(35.1%) | 28<br>(18.9%) | - |
| *Rep. 3* | 147 | 53<br>(36.1%) | 22<br>(15.0%) | 51<br>(34.7%) | 21<br>(14.3%) | - |

**Extended Data fig. 1:**
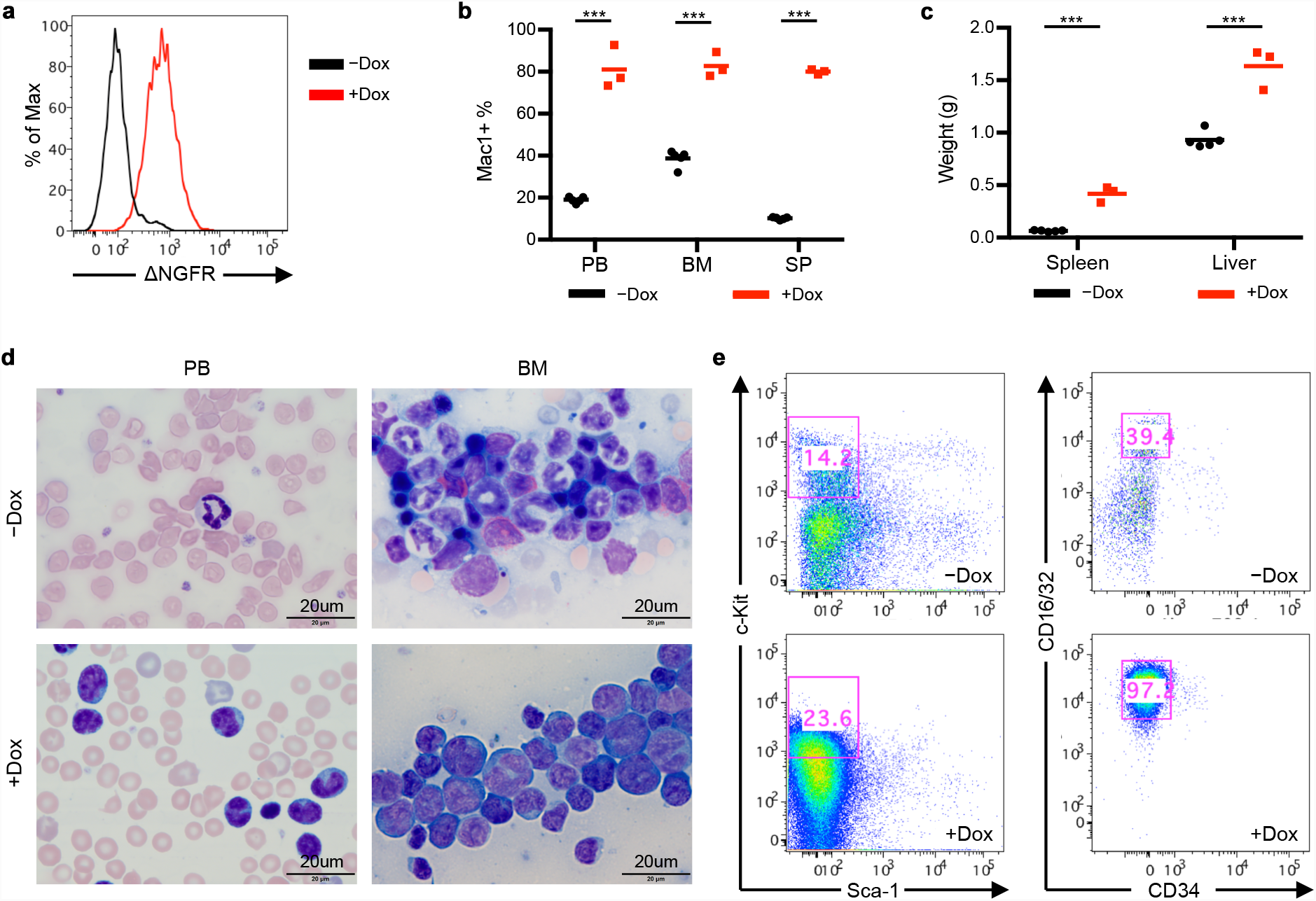
iMLL-AF9 GMPs support AML development *in vivo.* a, FACS analysis of NGFR expression in iMLL-AF9 GMPs following doxycycline (Dox) addition. Analysis was performed 24 hours after +/-Dox. b, Myeloid cell percentage in peripheral blood (PB), bone marrow (BM), and spleen (SP) of recipient mice transplanted with iMLL-AF9 GMP. Samples were collected and analyzed 7 weeks post transplantation. p < 0.001 for all comparisons. (n=5 for -Dox group, n=3 for +Dox group) c, Spleen and liver weights of recipient mice transplanted with iMLL-AF9 GMPs. Samples were collected and analyzed 7 weeks post transplantation, p < 0.001 for all comparisons. (n=5 for -Dox group, n=3 for +Dox group). Data from a repre sentative cohort is shown. d, Giemsa staining of representative peripheral blood (PB) smear and bone marrow (BM) smear from iMLL-AF9 GMP transplanted recipient mice 7 weeks post transplantation. Scale bar: 20 µm. e, FACS plots confirming the presence of L-GMPs in diseased mouse bone marrow.

**Extended Data fig. 2:**
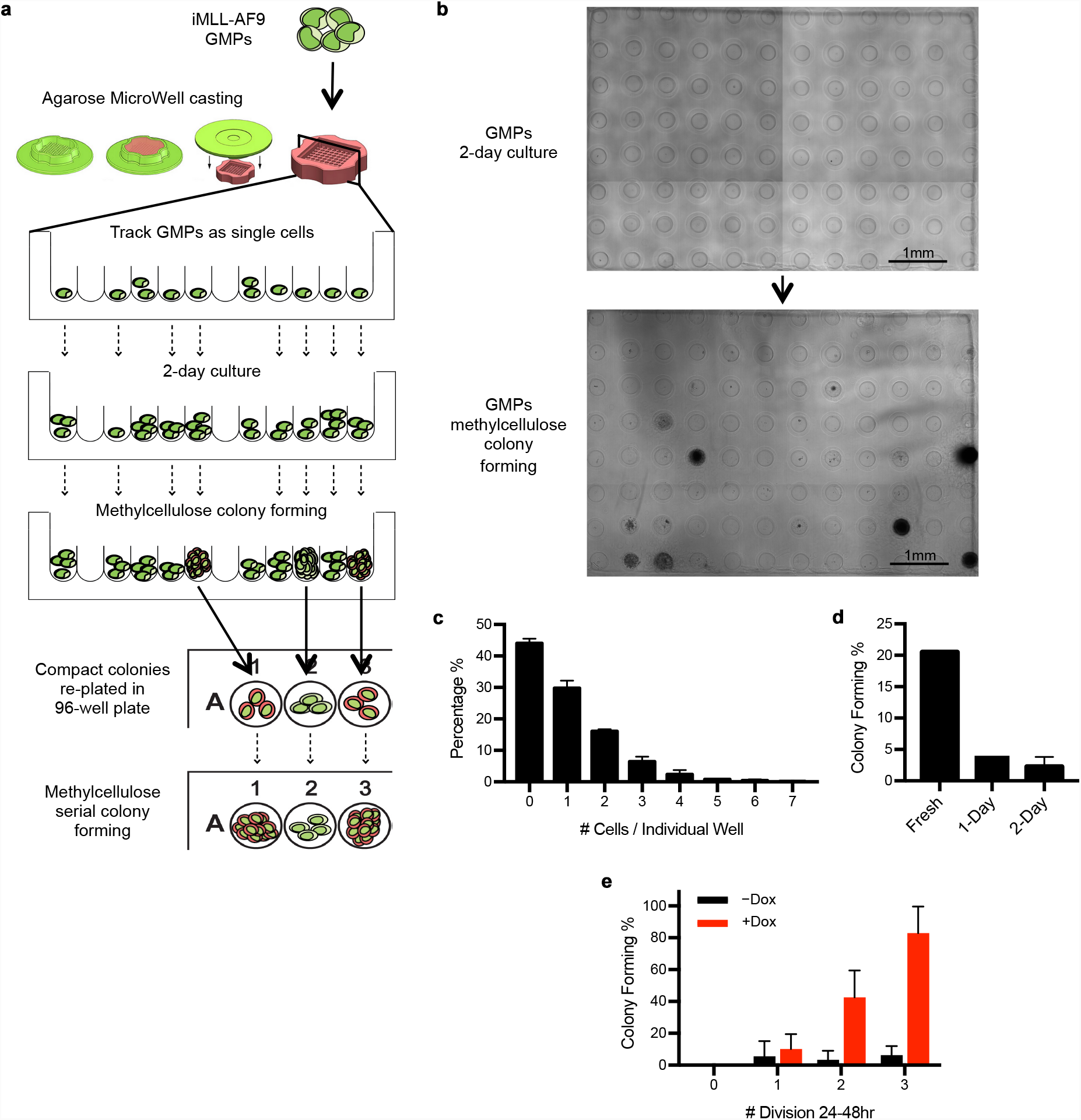
Permissiveness for MLL-AF9 mediated transformation is associated with the intrinsic cy cling rate of GMPs. a, Detailed workflow to track transformation from single iMLL-AF9 GMPs, as shown also in Fig. 1 d. A schematic cross-section view of the micro-wells is shown. Only the wells that contained single cells right after cell plating were continuously tracked. To confirm serial replating ability, colonies confined in each well were plucked and replated into single wells in a 96-well plate. The presence of compact colonies in 96-well plates was scored as successful replating events. b, Representative images of one micro-well gel during liquid culture (top) and following methylcellulose colony formation (bottom). Cells shown in these images were GMPs transduced with a lentivirus expressing MLL-AF9, which shares the same coding sequence for the iMLL-AF9 allele. Images were taken with a 4X objective, and stitched to show the entire micro-wells. Scale bar: 1 mm. c, Proper cell plating conditions for obtaining large numbers of wells that contain single cells. Typical cell number distri bution in individual wells when 75ul of cells resuspended at 2,500 cells/ml were loaded into one micro-well unit. Shown cell number distribution is a summary of four independent experiments, with 4-5 micro-well units analyzed for each ex periment. d, Primary colony forming efficiency in methylcellulose by single iMLL-AF9 GMPs, when they were freshly isolated or following one or two days of culture. e, Primary colony forming efficiency by iMLL-AF9 GMPs of different cycling rates, in the absence or presence of Dox. Cycling rates were defined by number of cell divisions during 24-48hr *in vitro* culture. Note this cycling rate was no longer the intrinsic cycle rate, as shown in Fig. 2d.

**Extended Data fig. 3:**
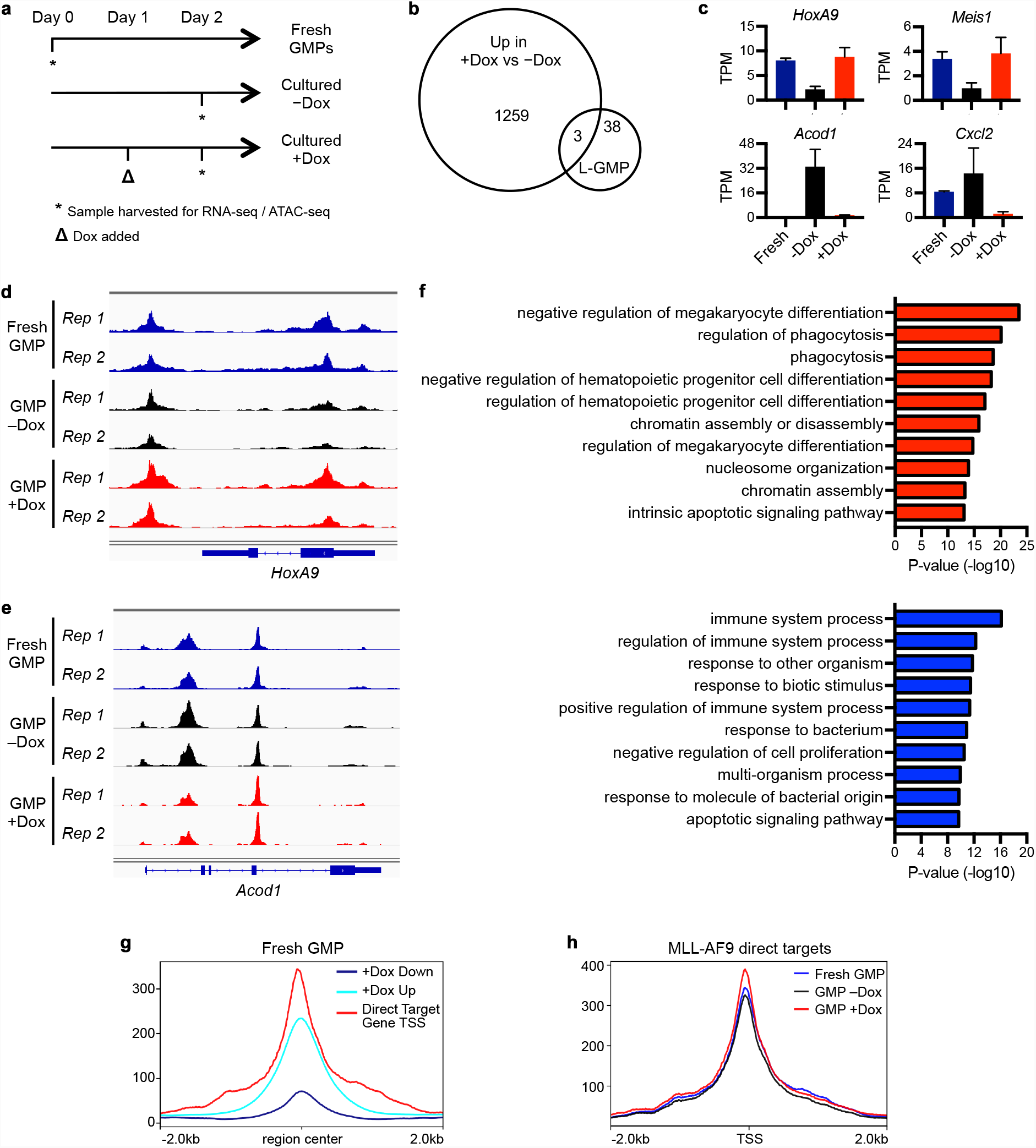
MLL-AF9 helps to sustain the already-existing gene expression program in GMPs. a, Schematic showing the timeline for iMLL-AF9 GMP treatment with Dox and sample harvest, for both RNA-Seq and ATAC-Seq analyses. b, Venn diagram showing minimal overlap between the Dox up-regulated DEGs as defined in Fig. 3a and the previously defined stem cell signature in L-GMPSB. c, Expression level changes of representative up-/down-regulated DEGs as defined in Fig. 3a. d, Representative ATAC-seq tracks, with *HoxA9* genomic regions as an example, showing increased chromatin accessi bility in +Dox versus -Dox iMLL-AF9 GMPs. e, Representative ATAC-seq tracks, with *Acod1* genomic regions as an example, showing decreased chromatin accessi bility in +Dox versus -Dox iMLL-AF9 GMPs. f, GO analysis of the differential ATAC-Seq peaks from iMLL-AF9 GMPs treated +/-Dox for 24 hours, as defined in Fig. 3f. The top 10 biological process terms up-regulated (red bars) and down-regulated (blue bars) in +Dox GMPs are shown. g, The direct targets of MLL-AF9 (red line, defined by MLL-AF9 CHIP-Seq^22^) were highly accessible in fresh GMPs, beyond those more accessible regions caused by Dox induction, as defined in Fig. 3f. h, 24 hours of Dox induction (red line) promoted further opening of the genomic regions directly bound by MLL-AF9 (de fined by MLL-AF9 CHIP-Seq 22}, which display significant accessibility in fresh GMPs (blue line). 4

**Extended Data fig. 4:**
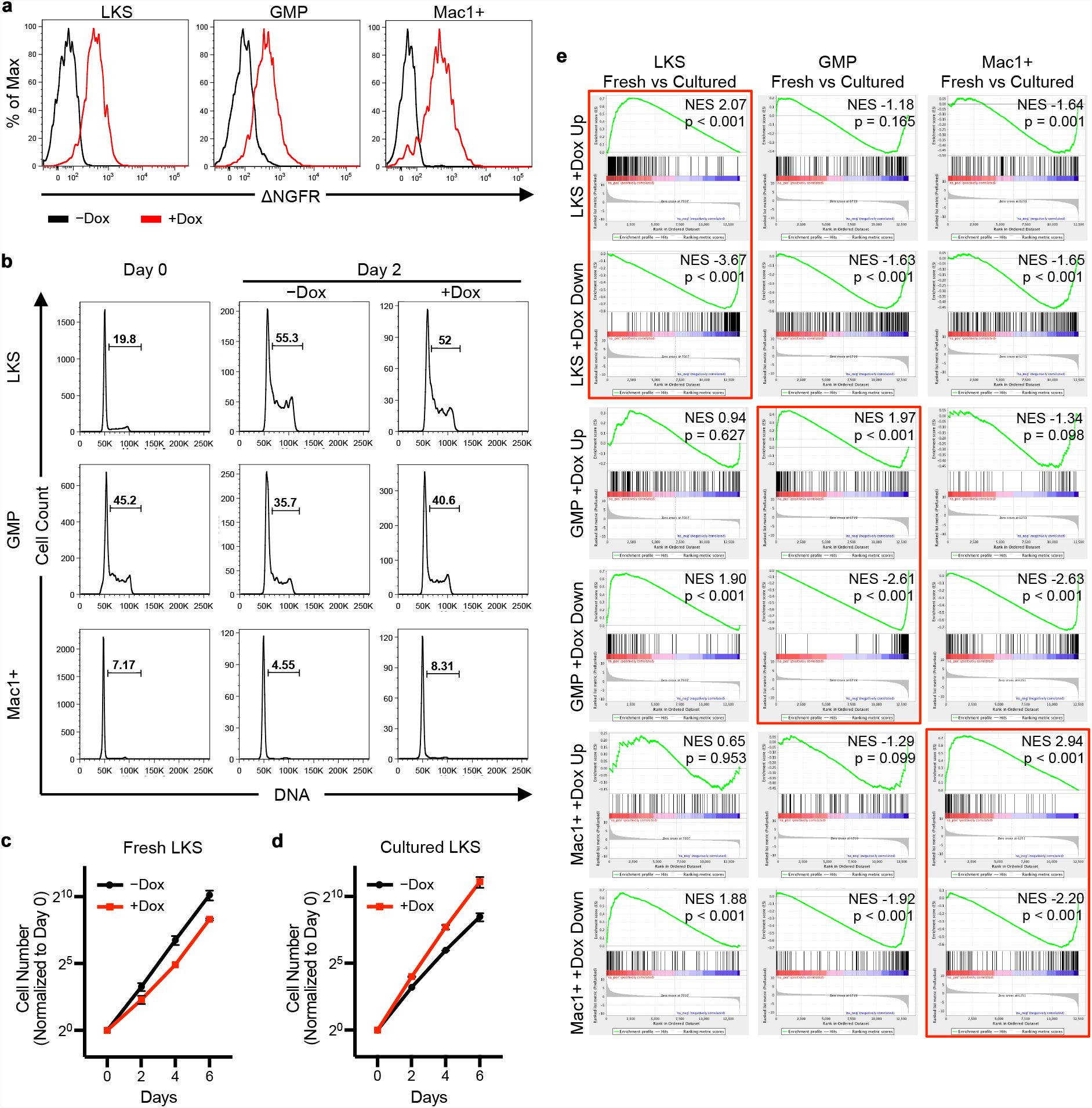
The primary gene expressionchanges to MLL-AF9 induction iscell-context dependent. a, FACS plots of hNGFR staining in LKS, GMP and Mac1+ cells, confirming MLL-AF9 induction in all cell types. b, FACS plots of Hoechst staining of DNA content in specified fresh cell types, or after 2 days of culture +/-Dox. Gates denote the frequency of cells in S/G2/M phases of the cell cycle. c, Proliferation response to MLL-AF9 induction in fresh LKS cells. Dox was added to fresh LKS at Day 0. d, Proliferation response to MLL-AF9 induction in cultured LKS cells. Dox was added after 4 days. x-axis denotes the time (Days) after Dox addition. Note the relative position of the black line and red line is now switched as compared to those inc. e, The Dox-induced DEGs, similar to those defined in Fig. 4d but at p<0.01, were queried against the gene expression changes occurred during culture across all three cell types by GSEA. For GMPs, only the top 200 DEGs were used. Note the DEGs were only enriched for their own respective cell type, but not across the other two cell types. DEGs against their respective cell types were highlighted by red box, and are the same as shown in Fig. 4g.

**Extended Data fig. 5:**
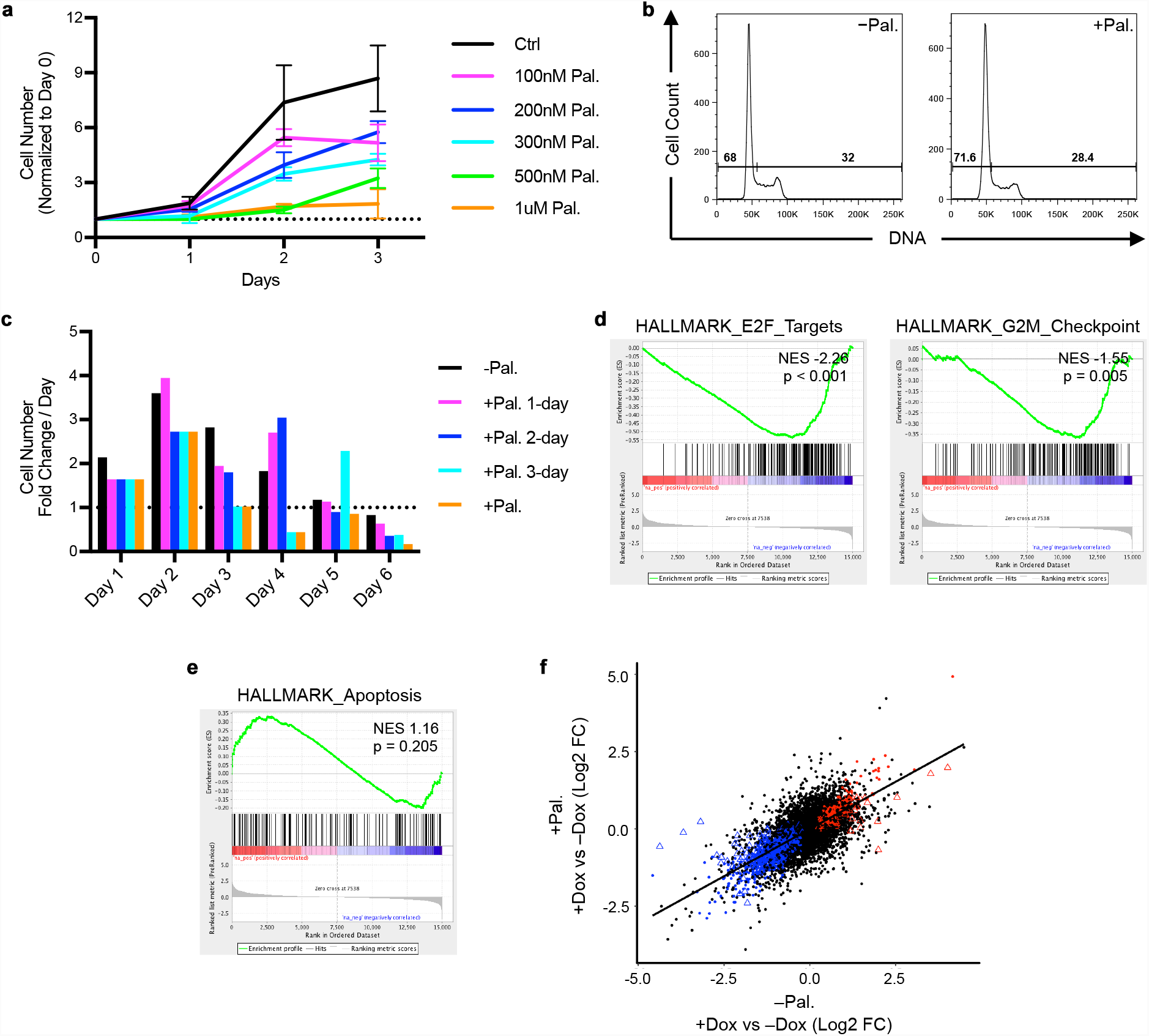
Modification of the initial cell state by cell cycle inhibition mitigates MLL-AF9 mediated changes in gene expression. a, GMP proliferation rates in the presence of different concentration of palbociclib. Proliferation was determined by scor ing the number of viable cells in each condition at daily intervals. Note the minimal reduction in cell numbers at 500 nM (green line). b, FACS plots of Hoechst staining of DNA content of GMPs following a 2-day culture in the presence of 500nM palbo cilib. Gates denote the frequency of cells in S/G2/M phases of the cell cycle. c, The proliferation rates of GMPs treated with 500nM palbociclib for various durations. For all treated conditions, palbo ciclib was added on Day 0, and washed out following the indicated duration. Cell proliferation rate (y-axis) was mea sured as cell number normalized to that of the same condition on the previous day. Dotted line (y= 1) indicated no change in cell number during the indicated two consecutive days. Note the recovery of proliferation in all treated condi tions following palbociclib removal, except for when it was present throughout. d, GMPs treated with 500nM palbociclib for 2 days down-regulated cell cycle pathways as determined by GSEA. e, GMPs treated with 500nM palbociclib for 2 days did not activate apoptotic pathways significantly as determined by GSEA. f, The modified gene expression response to MLL-AF9 induction (+/-Dox) by palbociclib (+ or -Pal.). Individual genes were plotted as single dots or triangles. Black line is the linear regression of the scatter plot, showing positive correla tion. Significantly up-regulated and down-regulated genes are marked in red and blue, respectively. Genes similarly changed by Dox irrespective of the presence of Pal. are denoted as dots, while those changed only in the absence of palbociclib are denoted as triangles.

**Extended Data fig. 6:**
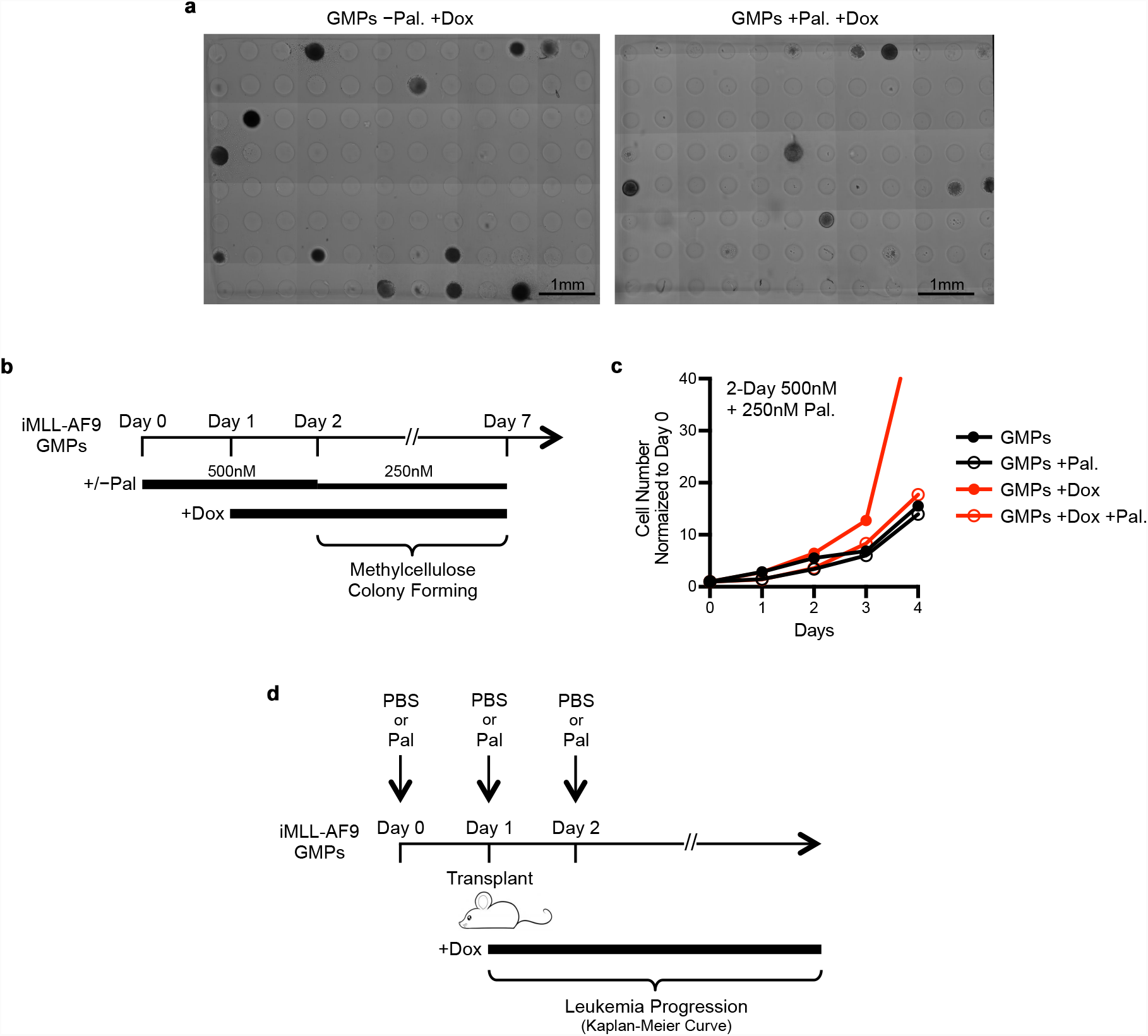
Modification of the initial cell state by mild cell cycle inhibition mitigates MLL-AF9 mediat ed transformation. a, Representative images of transformed colonies formed by iMLL-AF9 GMPs in micro-wells, in the absence or pres ence of palbociclib. Scale bar: 1 mm. b, Schematics illustrating the timeline for colony formation by bulked cultured iMLL-AF9 GMPs +/-Pal. c, The proliferation rates of bulk cultured iMLL-AF9 GMPs in various treatment conditions following the first 4 days of the schematics shown in c. d, Schematics illustrating the timeline for *in vivo* leukemogenesis by iMLL-AF9 GMPs, following the mild and temporary treatment by palbociclib. PBS was the vehicle control for palbociclib.

